# Decoding Internal Decision Making During Reverse Engineering Tasks

**DOI:** 10.1101/2023.10.10.561734

**Authors:** Brianna Marsh, Jocelyn Rego, Mia Matsuhisa Levy, Mitchell Peter Sayer, Alexander Waagen, Aidan Barbieux, Edward A Cranford, Donald F Morrison, Froylan Maldonado, Jeremy Phillip Johnson, Joseph DiVita, Jonathan Michael Buch, Tiffany Hyun-Jin Kim, Christian Lebiere, Sunny Fugate, Rajan Bhattacharyya

## Abstract

Neural decoding is often limited to tasks with known stimuli and limited response options . Real world tasks, however, are often completely stimulus free with unconstrained user response possibilities. Real time decoding of internal decision making would allow for more complex and interactive Huma Machine Teaming in a way that is not currently possible. To address this problem, we present here a novel method of decoding moments of recognition and their associated internal value judgments in the context of highly complex software reverse engineering tasks. This is done through a combination of P300 detection (a neural marker of recognition) and the Engagement Index (a ratio of neural band powers) to determine whether an item has been identified as relevant to the task (to be further explored) or irrelevant to the task (to be quickly ignored). Artificial neural networks were trained to identify P300s in each subject during the reverse engineering tasks. Dimensionality reduction of neural data during the tasks showed the existence of separately clustering subgroups of P300s with differences in Engagement Index. Subgroups of P300s differentiated by Engagement were further verified as distinct groupings with pupil dilation and user behavior metrics. This decoded information could be used to aid in the reverse engineering process via cognitive offloading of the user’s own decision making on to the visual interface in a completely automated and personalized fashion. This represents a significant advance in domain of real-time neural decoding, and opens up many further possibilities for usage in a broad range of intelligent human systems integration applications.

## INTRODUCTION

Real-world, complex, and unstructured tasks are rarely tackled in brain computer interface (BCI) research. Though challenging, the development of BCIs that can identify cognitive states in a complex context is an area of research that should not be left unexplored. This project aims at understanding the cognitive processes of reverse engineers (REs) as they perform vulnerability analysis on a binary file.

A key strategy of expert RE’s is to very quickly exclude large portions of the code as irrelevant and focus only on a much smaller subset of potentially vulnerable code (Mantovani, 2022). In this paper, we propose a novel combination of neural signals that can be used to identify and differentiate moments of recognizing relevance or irrelevance during a complex task such as reverse engineering. Identifying the moment of recognition is first done with an event related potential called the P300. This is a positive deflection in the EEG signal which can be detected approximately 300 ms after a surprising, rare, or particularly attended to event (Linden, 2005). In addition to the classic P300, a variety of “No-Go” P300s have been noted in the literature - these are P300s that occur during common or non-target stimuli (Polich, 2007). Importantly, identification of No-Go P300s is primarily done by having known differential stimuli or post-hoc analysis of differences in latency, amplitude, or lateralization of the signal. However, these methods are not sufficient for complex, stimulus-free, real time decoding: additional neural information is needed to make the distinction.

EEG frequency bands are typically defined as delta (1-4 Hz), theta (4-8 Hz), alpha (8-12 Hz), beta (12-30 Hz), and gamma (30-100 Hz), each of which has been associated with a range of cognitive states. The Engagement Index (EI), which combines beta, theta, and alpha band powers, has been used to quantify mental attention (MacLean, 2012) and alertness (Freeman, 1999). We here show that P300s in this task naturally cluster into separate groups, differentiable via the Engagement Index. Distinction between these two different types of neural events is then validated with user activity and pupil data. In combination, these two key metrics can decode Go vs. No Go P300 events in real time and give insight into an RE’s internal decision making in a completely automated fashion.

These insights could be used to augment a task interface, offloading retention of knowledge from human to machine. These two different types of moments of recognition may have many similarities, but require opposite interventions - bring attention to an area if it is relevant, or completely discard if it is not. Importantly, being able to modify the task interface to reflect these internal judgments has the potential to provide significant cognitive offloading to the user in providing a visual display of their intuitions, allowing them to further narrow down relevant subsections of code and work in a more efficient manner. Our method represents a major advance in real-time neural decoding and intelligent human systems integration as field.

## METHODS

Our subjects consisted of nine reverse engineers (RE’s) from HRL Laboratories and the Naval Information Warfare Center. These RE’s conduct their tasks in a platform called Ghidra, a reverse engineering tool developed by the National Security Agency (2019). In our experiment, participants first completed a number of unrelated pre-tasks as neurophysiological baselines. They were then given a series of 6 reverse engineering problems from the DARPA CHESS Challenge set (Kerley, 2022) and asked to identify the vulnerability in a given section of code (if one exists). During this process we collected recordings of their neural activity via EEG, eye tracking data, and actions in the tool interface (clicks, scrolls, tool changes, etc.). IRB approval was attained for this experiment; participants were compensated according to their normal hourly rates..

### Pre-tasks

Prior to the main experiment, participants completed an Auditory Oddball task. Here, participants sat with their eyes open while intermittent tones were played. Participants were instructed to press a key when they hear a high tone (Target), and ignore low tones (Distractor).These 2 stimulic lassically evoke Go and No Go P300 waveforms. Data was also collected while participants sat quietly with eyes open and no other stimuli occurring for several minutes.

### User behavior collection

The behaviors of reverse engineers were recorded as they interacted with the Ghidra interface with Swing event listeners and Java’s Abstract Window Toolkit implemented on Ghidra components. Behavioral data collected included clicks, mouse movements, scrolls, and keyboard inputs.

### Eye tracking

The Smart Eye AI-X eye tracking device was used to track eye fixations and movement on the computer screen interface. This device also measured pupil diameter of both the left and right eyes. As the experiments were performed in well lit rooms with constant luminance, external factors are expected to have minimal effect on pupil dilation. This method provided a 60 Hz sample rate.

### EEG

Recordings were made with a 32 channel Brain Vision ActiCHamp (subjects 1-3) or Neuroelectrics Enobio 32 (subjects 4-9) EEG cap with electrodes placed on the scalp according to the standard 10-20 scheme, at a sampling rate of 500 Hz. Signals were notch filtered at 60 Hz to remove power line artifacts and bandpass filtered between 1-128 Hz. Signals were then divided into 1 second epochs.

### P300 identification

Neural data from the Auditory Oddball test and eyes-open baseline period were used to train a machine learning model to identify P300 occurrences in neural data. Both Go and No Go P300s were labelled as a 1, while baseline data was labelled as a 0. Feedforward networks, described in figure 1, were trained via 5 fold cross validation. Each network was trained on 1 second windows of data from the FZ and CZ electrodes, flattened into a 1 dimensional vector, as the central and frontal midline regions are traditionally associated with the P300 signal (Uvais, 2018).

**Figure 1:**
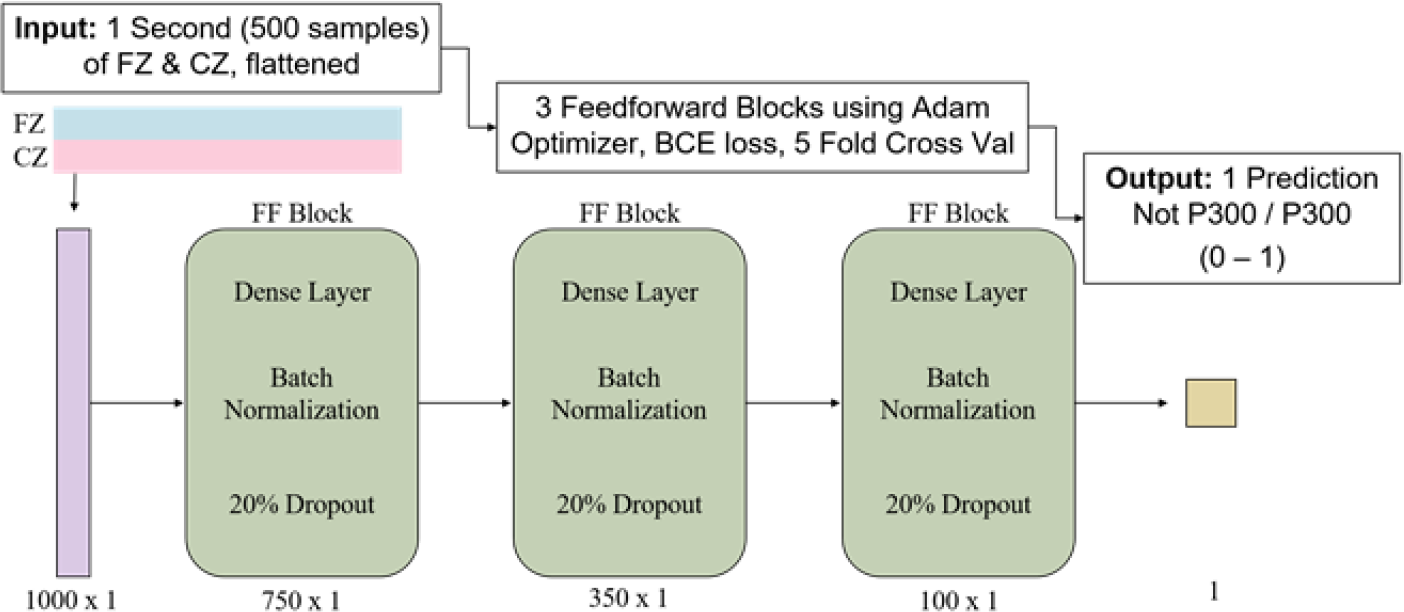
P300 Neural Network Architecture, displaying 3 feedforward (FF) blocks each containing a dense layer, batch normalization, and 20% dropout. The network is trained using 5-fold cross validation with binary cross entropy (BCE) loss and the Adam optimizer. Input is raw neural signal from midline electrodes FZ and CZ; Output node predicts if a one second epoch of EEG data contains a P300 or not.

### Band powers

The decomposition of EEG signals into frequency bands is here achieved using Welch’s method (Welch, 1967), and averaged across all scalp electrodes. The frequency bands were computed as delta (1-4 Hz), theta (4-8 Hz), alpha (8-12 Hz), beta (12-30 Hz), and gamma (30-100 Hz). Engagement Index (EI) is calculated as beta band power divided by the sum of the theta and alpha band powers for each one second epoch during the experiment.

### Dimensionality reduction

For each one second time point in the reverse engineers tasks, we created a feature vector that contained the raw P300 prediction as well as the average power in the 5 different frequency bands in each of the 32 EEG channels. We then employed UMAP (Uniform Manifold Approximation and Projection) (McInnes, 2019) to project these 161 dimensional vectors onto a 2 dimensional space

### Statistical analysis

Statistical analysis was performed to determine if there were significant differences between the High and Low Engagement P300 Points. For pupil diameter and standard deviation, a mixed effects model with subject ID as the grouping variable was employed to account for within-subject variation (Bates, 2014). Otherwise, T-tests were used. All tests required p < 0.05 to be considered statistically significant.

## RESULTS

### P300

All 9 subjects showed strong responses to both the Target high tones and Distractor low tones in the Auditory Oddball task, reflecting both “Go” and “No Go” type P300s. Representative average neural signatures from the midline electrodes for all stimuli can be seen in figure 2.

**Figure 2:**
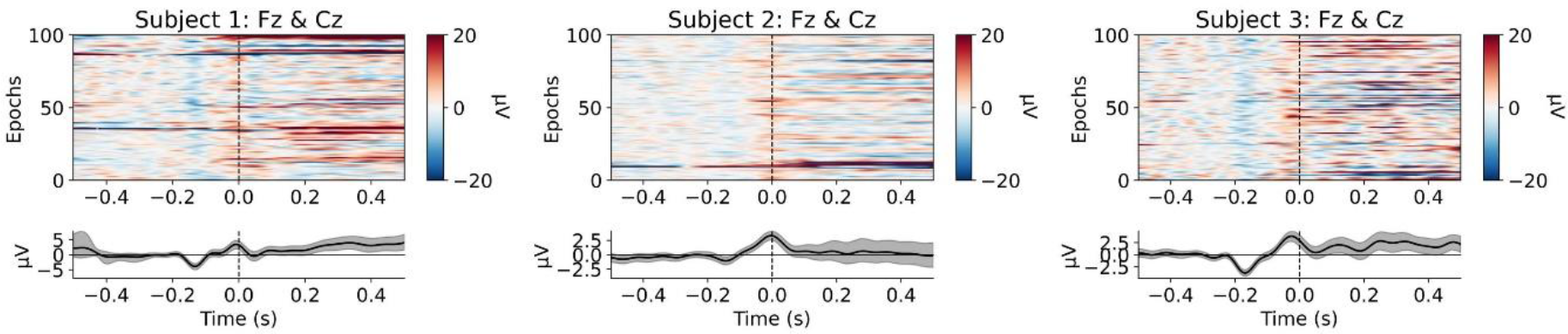
Sample waveforms for 3 subjects from the Auditory Oddball task (including both Go and No-Go P300s). Top: All trials (y axis) by time (x axis), where time 0 is the expected P300 peak time 300 ms after the stimulus. Bottom: Average waveform across all trials.

Across all 9 subjects, trained artificial neural networks showed extremely consistent results with an average validation accuracy of 96.36% (±1.75) across all subjects (Table 1). These networks were trained individually per subject, then applied to each subject’s respective neural data during the reverse engineering tasks, broken into 1 second time windows. The networks predicted an average of 13.72% of all time windows to contain a P300, with an average of 6.7 unique events per minute. Continuously identified P300 predictions lasted approximately 1.2 seconds, as would be expected for a short physiological signal.

**Table 1.** Top - Oddball validation accuracy using five-fold cross validation. On average, networks achieve 96.36% accuracy ±1.75% (standard deviation), showing consistency across both training folds and subjects. Bottom – Reverse Engineering task results. On average, subjects correctly answered 2.33 out of 6 tasks (39%). An average of 6.7 P300 events were detected per minute during the tasks, with a sustained P300 detection duration of 1.21 seconds. Across all time in the tasks, 13% of time points were predicted to contain a P300; of all detected P300 time points, 15.91% showed High Engagement.

| P300 Neural Network Accuracy |  |  |  |  |  |
| --- | --- | --- | --- | --- | --- |
|  | Fold 1 | Fold 2 | Fold 3 | Fold 4 | Fold 5 |
| Average | 95.84 | 96.82 | 96.87 | 96.06 | 96.23 |
| Standard Deviation | 1.43 | 1.48 | 1.14 | 2.67 | 1.80 |
| Reverse Engineering Task Results |  |  |  |  |  |
|  | # Correct | Events/Min | Duration (s) | % Time | % High P300 |
| Average | 2.33 | 6.70 | 1.21 | 13.72 | 15.91 |
| Standard Deviation | 0.87 | 2.05 | 0.09 | 5.33 | 8.46 |

### Engagement Index

Engagement Index was calculated as the below ratio of neural band powers for each electrode per second per subject.

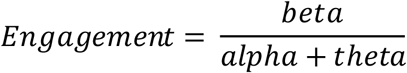

A threshold for High Engagement was set as 1 standard deviation above the mean Engagement from the first 10 minutes of the reverse engineering tasks; everything below this threshold was classified as Low Engagement. Using this threshold, less than 16% of P300s identified during the reverse engineering tasks show High Engagement (Table 1). This aligns with our expectations that the majority of the P300s are due to rejecting lines of code rather than finding something interesting (due to sparsity of true vulnerabilities).

### Dimensionality reduction

To find clusters of cognitively similar time points across the reverse engineering tasks, we created UMAP projections of the P300 prediction and average band powers (Alpha, Beta, Gamma, Delta, & Theta) across all electrodes at every time point for each subject. In all 9 subjects, we consistently found subgroups of P300 points that separate both from the control data and each other (figure 3). This implies that there are 2 distinct phenomena that are classified as a P300 by the network - presumably, putative Go and No Go P300s. These are hypothesized to represent either moments when the RE identifies a piece of code that is relevant to a vulnerability, or moments when they identify information that allows them to exclude the code and quickly move on.

**Figure 3:**
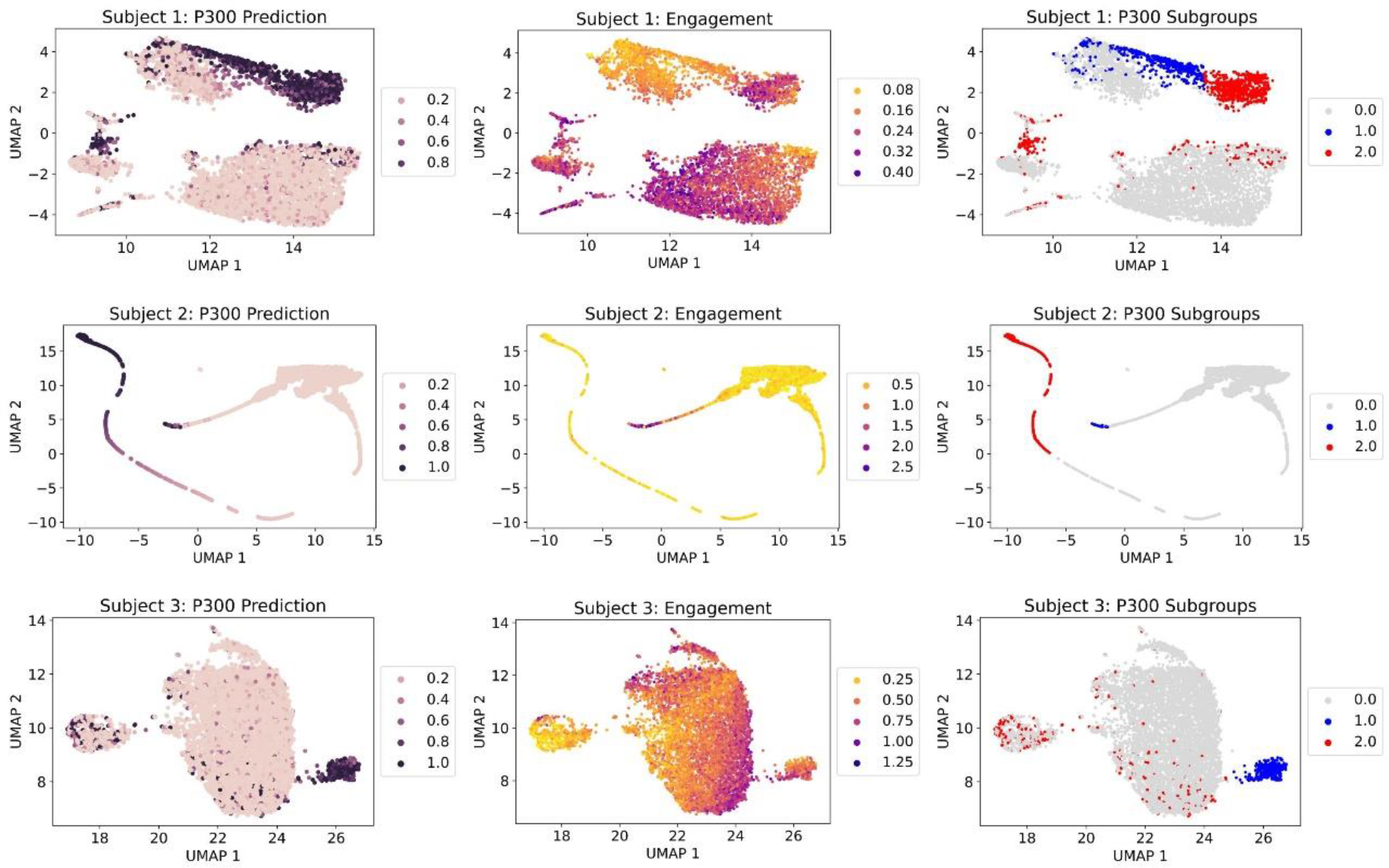
UMAP projection of the neural data of all time points for 3 subjects, colored by: P300 Prediction, Engagement Index (average over all channels), and Subgroup Labels. The Engagement Index is statistically significantly different between P300 Groups (blue and red dots) in an average of 22.89 channels (±8.84) out of 36 total channels per subject.

To test the hypothesis that the Engagement Index may explain differences between P300 subgroups within the UMAP project, we show the same projections with points colored by their calculated Engagement Index and their manually identified subgroup labels. Indeed, we see that there are visually obvious differences in Engagement between P300 subgroups. Statistical testing of the Engagement between the two P300 subgroups confirms that statistically significant differences exist across all subjects in an average of 22.89 (± 8.84) out of 36 total electrodes.

### Validation with pupil metrics

We then asked if the different groups of P300s could be further validated outside of neural data. Specifically, pupil dilation has been shown to be closely tied to Noradrenaline activity in the Locus Coeruleus region of the brain and can be taken as a proxy for activation of the sympathetic nervous system (Eckstein, 2017). Due to this, we looked at changes in pupil diameter as a secondary indication of differing underlying neural states between the two identified types of P300s (figure 4). The normalized (per participant) mean and standard deviation of the averaged pupil diameter per second was tested for significant differences between High Engagement P300 points and Low Engagement P300 points across all subjects using a multilevel model; both were determined to be statistically significantly different between groups (Mean: p = 0.003, SD: p < 0.000).

**Figure 4:**
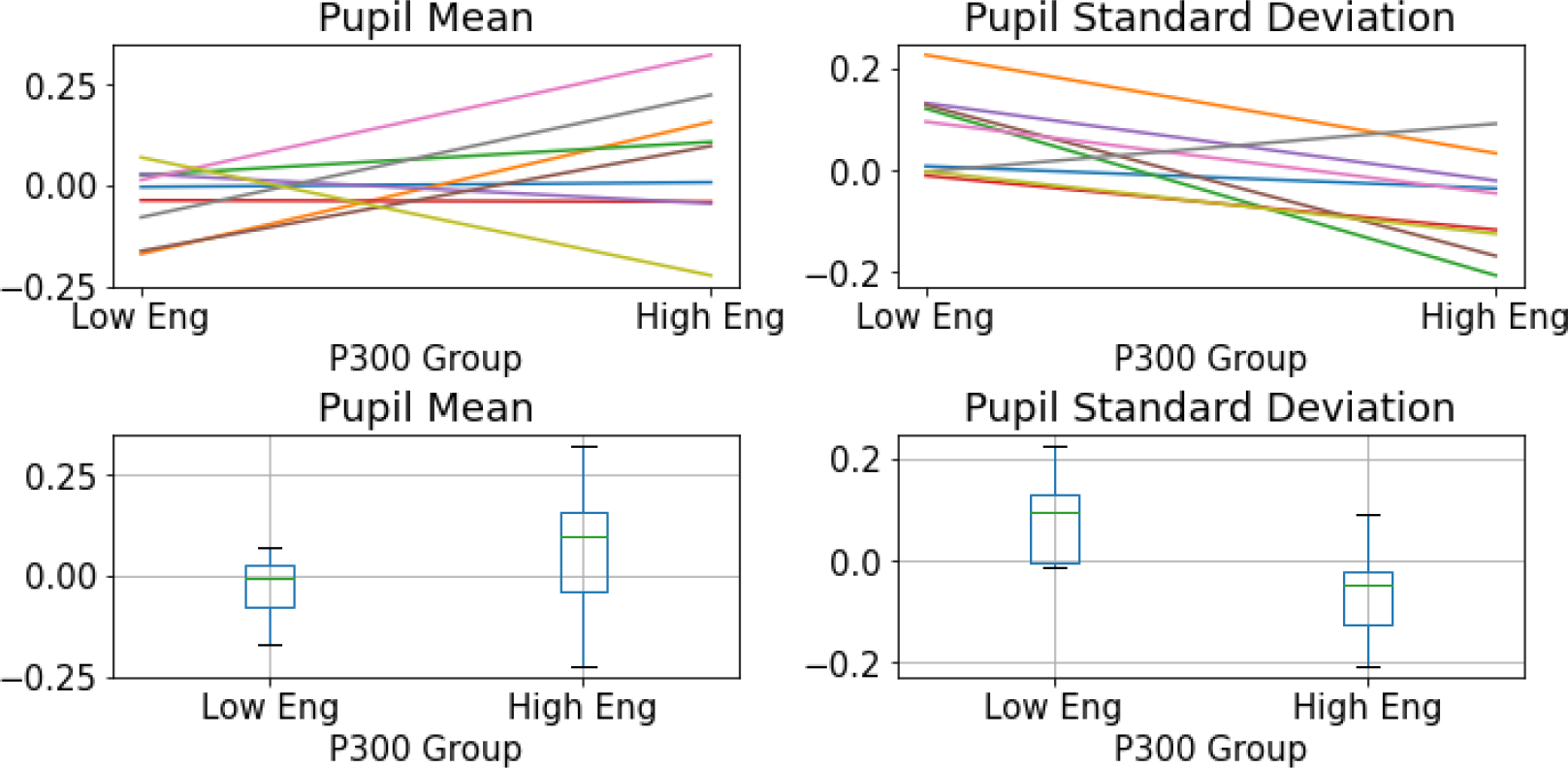
Normalized pupil diameter means (left) and standard deviations (right), averaged per participant per P300 Group (Low vs High Engagement). Top plots show regression lines per subject from the Mixed Effects Model to account for individual variation. High Engagement P300 points show a higher pupil diameter mean, and lower standard deviation (Mean: p = 0.003, SD: p < 0.000, Mixed Effects Model).

While differences existed between subjects, the majority of subjects showed Low Engagement P300 points with a lower mean and higher standard deviation; this could indicate they are more relaxed and having normal pupil fluctuations. High Engagement P300 points then showed higher mean and lower standard deviation, indicating their pupils are steadily widened as they potentially have a moment of insight.

### Validation with user actions

We further validated our use of P300s and Engagement by analyzing the behavior of reverse engineers. The primary behavior of interest is a click followed by scrolling; this sequence was identified as highly likely to occur when reverse engineers find a potential vulnerability in code and search for further information to confirm or reject their hypothesis.

To test this hypothesis, we calculated the percentage of all High (and Low) Engagement P300s that occurred within 2 seconds before this sequence. We found that a significantly higher percentage of High Engagement P300s occur prior to a click-scroll, as compared to the percentage of Low Engagement P300s (p =0.0077, T-test). On average, subjects had 2.84% of all of their High Engagement P300s occur before this sequence, but only 0.22% of Low Engagement P300s (figure 5).

**Figure 5:**
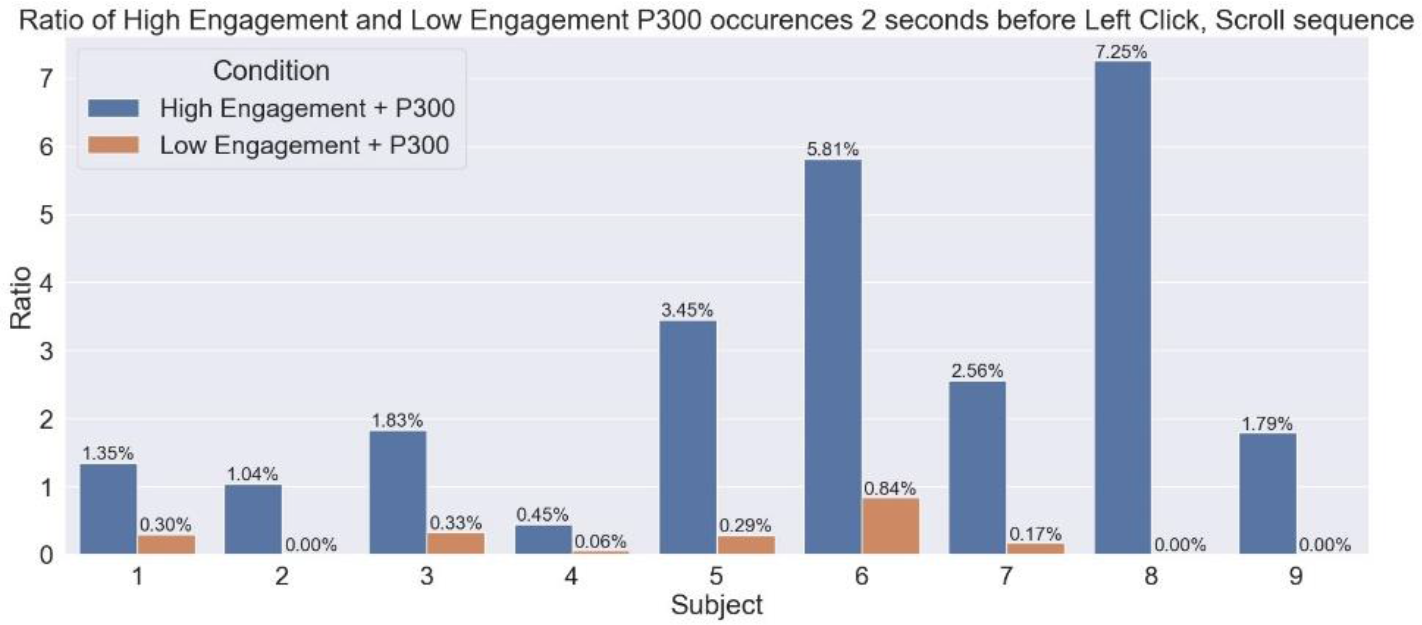
Percentage of all High (and Low) Engagement P300s that occur in the 2 seconds prior to a click-scroll sequence for each subject during reverse engineering tasks. On average, 2.84% (± 2.29) of all High Engagement P300s occurred before this sequence; the same is true for 0.22% (±0.27) of all Low Engagement P300s. This difference is statistically significantly different (p = 0.0077, T-test)

## DISCUSSION

We here propose a method of neural decoding based on two well established metrics - the P300 and the Engagement Index - in novel combination to differentiate Go and No-Go type P300s in real time. We validated our approach with neural data dimensionality reduction, pupil dilation, and behavioral data. This disambiguation of neural signals can be used to aid human users in complex, real world tasks; specifically, those that require human expertise in parsing large amounts of data to find sparse items of importance. This advance in Human-Machine Teaming allows for significant cognitive offloading, as the task interface can be automatically updated to reflect and track the user’s own internal decision making. This plays directly to the strengths of both parties - human task expertise and machine memory capacity.

For use in software reverse engineering, code lines of interest could be automatically highlighted in yellow while irrelevant lines could be automatically greyed out. Beyond reverse engineering, consider a lawyer who needs to parse large documents of legal text in search of scarce laws and precedents relevant to his client’s case. Medical applications could further span from aiding doctors in analyzing patient test results and medical scans, to augmenting classic Brain Computer Interface approaches for communication.

The primary limitation of this study is the lack of definitive ground truth during the reverse engineering tasks. Explicitly asking the RE to press a different button each time one of these events occurs is likely to be unreliable, distracting, and deeply entangle the signals of interest with artifacts from the movement itself and the additional cognitive processing of deciding to push a button. Measures of how helpful and intuitive our proposed intervention is can come only from user feedback in a real-time field test. Fine-tuning this tool for professional use will require extensive instrumentation and engineering of the task interface, and is beyond the scope of this paper. Here, we emphasize the scientific advances before the engineering of implementation.

Moving forward, we hope to explore more joint measures of neural activity to decode cognitive states. For example, eye gaze entropy has also been shown to be a valid metric for fatigue (Shiferaw, 2018) and could be used in combination with alpha band power (a robust metric for alertness (Knyazev, 2016)) as a metric of motivation and focus. Taken together, these could give a strong indication of when the user may require a break in order to continue performing efficiently. We are further exploring joint metrics for broad cognitive states such as stress, cognitive load, insight, and exploration vs. exploitation.

This metric is a starting point from which we hope to gain further meaningful insight into the cognitive states of human users during complex, real-world tasks. The ability to identify different types of P300s in real time without known external stimuli further presents the opportunity for an incredibly wide array of applications, and we hope can have far reaching impacts in the future of neural decoding and intelligent human machine teaming.

## ACKNOWLEDGMENT

This material is based upon work supported by the Defense Advanced Research Agency (DARPA) under Contract No. HR0011-20-C-0141. Any opinions, findings and conclusions or recommendations expressed in this material are those of the author(s) and do not necessarily reflect the views of the Defense Advanced Research Projects Agency (DARPA).

## REFERENCES

Bates, D. et al. (2014). “Fitting Linear Mixed-Effects Models using lme4”. In: arXiv: 1406.5823

Eckstein, M. K. et al. (2017). “Beyond eye gaze: What else can eyetracking reveal about cognition and cognitive development?” In: Developmental Cognitive Neuroscience 25, pp. 69–91

Freeman, F. G. et al. (1999). “Evaluation of an adaptive automation system using three EEG indices with a visual tracking task”. In: Biological psychology 50.1, pp. 61–76.

Kerley, B. (2022). CHESS ACES. https://github.com/cromulencellc/chess-aces

Knyazev, G.G., Savostyanov, A.N., and Levin, E.A. (2006). “Alpha synchronization and anxiety: implications for inhibition vs. alertness hypotheses”. In: Int J Psychophysiol 59.2, pp. 151–158.

Linden, D.E. (2005). “The p300: where in the brain is it produced and what does it tell us?” In:Neuroscientist 11.6, pp. 563–576.

MacLean, M. H., Arnell, K. M. and Cote, K. A. (2012). “Resting EEG in alpha and beta bands predicts individual differences in attentional blink magnitude”. In: Brain and Cognition 78.3, pp. 218–229. ISSN: 0278-2626.

Mantovani, A. et al. (2022). “RE-Mind: a First Look Inside the Mind of a Reverse Engineer”. In: 31^st^ USENIX Security Symposium.

McInnes, L. and Healy, J. (2018). “UMAP: Uniform Manifold Approximation and Projection for Dimension Reduction”. In: ArXiv abs/1802.03426.

National Security Agency, US (2019). “Ghidra Software Reverse Engineering Framework”. In:361 https://github.com/NationalSecurityAgency/ghidra.

Polich, J. (2007). “Updating P300: An Integrative Theory of P3a and P3b”. In: Clinical Neurophysiology 118, pp. 2128–2148.

Shiferaw, B. A., Downey, L.A., and Westlake, J. et al. (2018). “Stationary gaze entropy predicts lane departure events in sleep-deprived drivers.” In: Sci Rep 8.2220.

Stikic, M et al. (2014). “Modeling temporal sequences of cognitive state changes based on a combination of EEG-engagement, EEG-workload, and heart rate metrics”. In: Frontiers in Neuroscience 8. DOI: 10.3389/fnins.2014.00342.

Uvais, N.A. et al. (2018). “Auditory P300 event-related potential: Normative data in the Indian population.” In: Neurol India 66.1, pp. 176–180.

Welch, P. D. (1967). “The Use of the Fast Fourier Transform for the Estimation of Power Spectra: A Method Based on Time Averaging Over Short, Modified Periodograms.” In: IEEE Transactions on Audio and Electroacoustics AU-15, pp. 70–73.

